# scLKME: A Landmark-based Approach for Generating Multi-cellular Sample Embeddings from Single-cell Data

**DOI:** 10.1101/2023.11.13.566846

**Authors:** Haidong Yi, Natalie Stanley

## Abstract

Single-cell technologies enable high-dimensional profiling of individual cells, therefore offering profound insights into subtle variation between specialized cell-types. However, translating the multitude of nuanced cellular profiles into meaningful per-sample representations is challenging due to heterogeneous cellular composition across individual profiled samples. To compute informative per-sample representations, we developed scLKME, a novel approach that uses a landmark-based kernel mean embedding method to convert multi-sample single-cell data into compact per-sample embeddings. Treating each sample as a distribution over cells, scLKME identifies landmarks across samples and maps these distributions into a reproducing kernel Hilbert space. Overall, scLKME outperforms state-of-the-art techniques in robustness, efficiency, accuracy, and practical usefulness of sample embeddings. Its application on a CyTOF dataset profiling immune responses in preterm birth highlighted its capacity to accurately identify patient-specific variations correlating with gestational age, suggesting broad applicability to multi-sample single-cell datasets with complex experimental designs. scLKME is available as an open-sourced python package at https://github.com/CompCy-lab/scLKME.

## 1 Introduction

Single-cell technologies have enabled detailed insights into cellular states using high-dimensional gene or protein measurements. Numerous tools have emerged to make sense of these data, based on clustering [1–5], visualization [6–10], and trajectory inference [11–15]. While these methods offer deep insights into variation between cells, there is a gap in computing per-sample representations that reflect between-sample variation. To understand how per-sample cellular heterogeneity drives per-sample experimental or phenotypic outcomes, a key task is to generate per-sample *embeddings*, in which each sample is summarized based on all profiled cells. Unlike single-cell embeddings which reflect within-sample cellular variation, sample embeddings prioritize the aggregated characteristics of cellular abundances enabling nuanced per-sample understandings [16, 17].

Generating sample-level representations involves multiple challenges. First, in addition to within-sample cellular variation across all measured features, samples vary based on their cellular heterogeneity patterns between phenotypes or conditions. Any algorithm to encode each sample must preserve both sources of variation and be invariant to the order in which cells were profiled [16, 18]. To address these challenges, clustering-based [17, 19, 20] and pooling-based methods [16, 21] have been developed to uncover and encode sample-level variation. Briefly, clustering-based methods partition cells into groups and define feature vectors based on cell-type abundances, and pooling-based methods encode samples by aggregating individual cell features through pre-defined pooling functions, such as max or mean pooling. Although demonstrated to be successful in many applications, these two types of methods still showed some limitations. For instance, clustering-based methods rely on accurate cell-type identification via clustering, inhibiting its full automation without human involvement. As for pooling-based methods, the pooling operation is a bottleneck, which may not be sufficient to extract features across multitude cells.

In this study, we introduce scLKME, a landmark-based approach that uses kernel mean embedding to compute vector representations for samples profiled with single-cell technologies (see Figure 1). By viewing each sample as a highdimensional distribution, scLKME first sketches or sub-selects a limited set of cells across samples as landmarks. To summarize the characteristics of persample distributions, scLKME maps them into a reproducing kernel Hilbert space (RKHS) using kernel mean embedding. The final embeddings are generated by evaluating these transformed distributions at the sampled landmarks, yielding a sample-by-landmark matrix. We demonstrated the effectiveness of scLKME in terms of accuracy, efficiency, and robustness by comparing its performance in capturing between sample-variation with existing methods [19–21]. In particular, scLKME’s utility was demonstrated on a CyTOF dataset of preterm births, and successfully generated per-sample encodings that captured inter-patient variation, such as, gestational age.

**Fig. 1.**
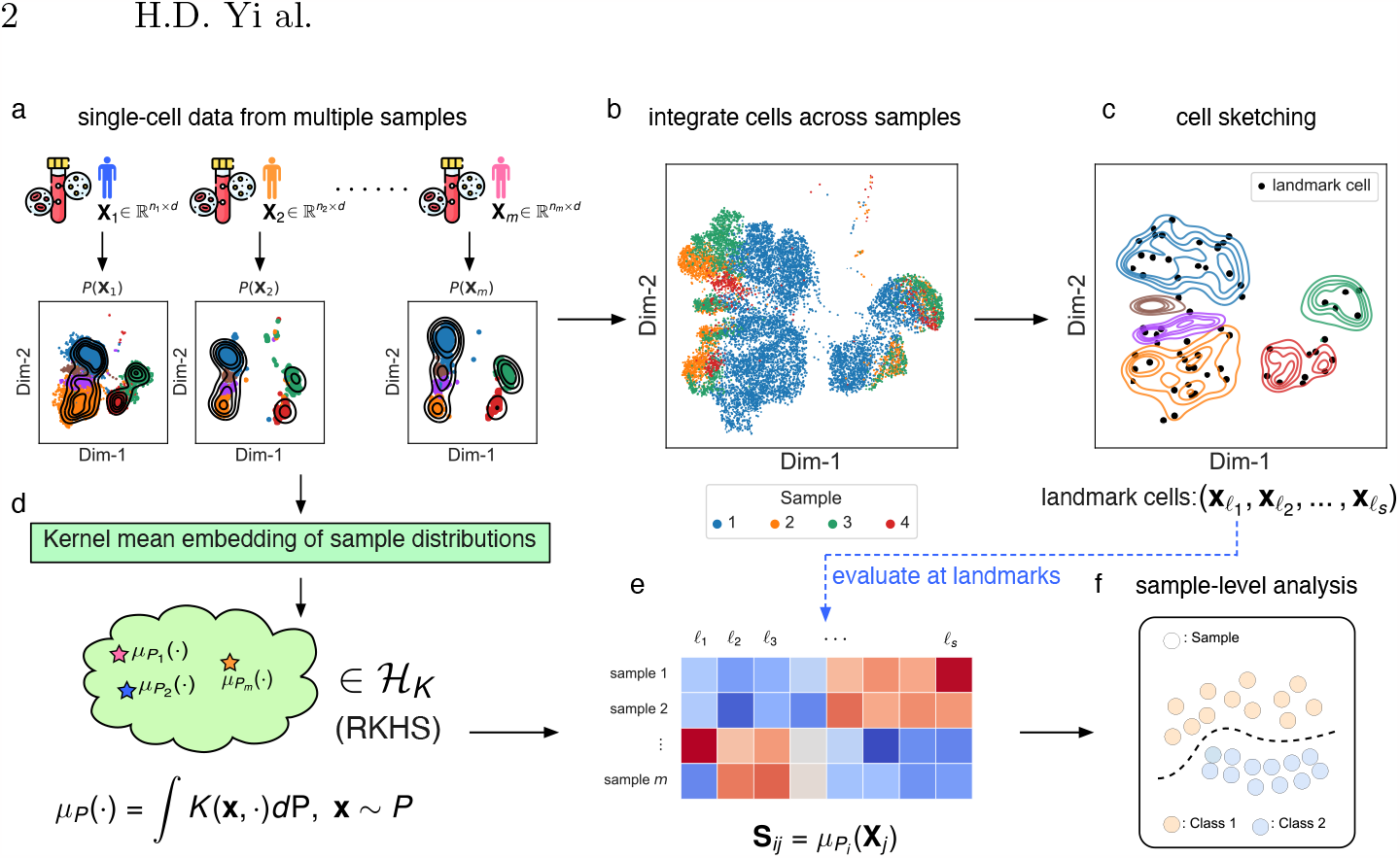
Flow diagram outlining the workflow of scLKME. **a**. Single-cell datasets are collected from multiple samples, where each sample is modeled as a high-dimensional distribution over profiled cells. **b**. Cells from distinct samples are aggregated, and then integrated into a shared space. **c**. The dataset is sketched by a small number of landmark cells (black points). **d**. Sample distributions are mapped into a reproducing kernel Hilbert space (RKHS) using kernel mean embedding. **e**. Sample embeddings are generated by evaluating the transformed distributions in the RKHS at the landmark cells. **f**. Sample-level downstream analysis is enabled based on sample embeddings.

## 2 Methods

### 2.1 Cell Sketching

The input to scLKME is an integrated dataset of cells across *m* total samples, denoted as 𝒟 = {**X**_1_, **X**_2_, …, **X**_*m*_}. Each sample 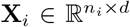 is a matrix encoding the expression of *d* gene or protein measurements across *n*_*i*_ cells. Associated with each sample are discrete or continuous response labels, denoted as {*y*_1_, *y*_2_, …, *y*_*m*_ }. As a preliminary step, scLKME creates a *sketch* or downsampled version of *s* cells across all samples. The sketched dataset is denoted as 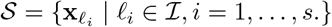. where *ℐ* is an indicator set containing indices of cells in the sketch. These sketched cells serve as landmarks, or anchors, in the dataset to ultimately compute per-sample kernel mean embeddings. Notably, the sketched version of *s* cells represents a small fraction of the dataset, with each 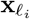originating from one of the samples. The parameter *s* can be tuned by users, allowing a trade-off between information content and computational complexity. scLKME offer three possible sketching methods, including, geometric sketching [22], kernel herding [23, 24], and simple random sampling without replacement.

### 2.2 Landmark-based Kernel Mean Embedding

For each sample, **X**_*i*_ where *i* = 1, …, *m*, we consider cells as independent and identically distributed (i.i.d.) variables from a distribution *P*_*i*_, with *i* = 1, …, *m*. Using kernel mean embedding (KME), we map these sample distributions into a reproducing kernel Hilbert space (RKHS). Formally, the KME of a probability measure *P* on domain *𝒳* is:

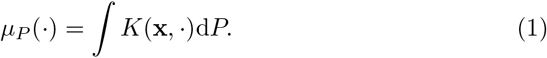

Here, the KME maps distribution *P* into RKHS, *ℋ* _*K*_, endowed by a reproducing kernel *K* : *𝒳 ×𝒳* ℝ [25]. The embedding *μ*_*P*_ (*·*) *∈ ℋ*_*K*_ serves as the representation for *P*, akin to a characteristic function. The utility of KME lies in its injective property. That is, for any two probability measures *P, Q* on *𝒳*, KME ensures that ∥*μ*_*P*_ − *μ*_*Q*_ ∥ *ℋ* = 0 if and only if *P* = *Q*. This is valid when the kernel is characteristic, a condition met by many common kernels, such as the Gaussian and Laplacian kernels [26, 27].

*Approximation of KME* Analytically calculating *μ*_*P*_ (*·*) is intractable due to unknown sample distributions. Therefore, we approximate *μ*_*P*_ (*·*) using the following unbiased empirical averaging estimator:

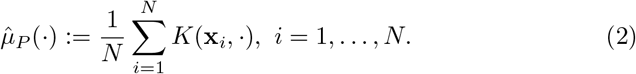

Here, **x**_*i*_, *i* = 1, …, *N* are i.i.d. samples from *P*. By the weak law of large numbers, this estimator (2) is guaranteed to converge to the true mean [28, 29].

KME derives various distribution discrepancy metrics, including *integral probability metric* (IPM) [30] and *maximum mean discrepancy* (MMD) [31]. While successful in statistical testing [31, 32] and machine learning [33, 34], their computational cost is high, especially for empirical distributions. For instance, given i.i.d. samples **X** = {**x**_1_, …, **x**_*N*_ }and **Y** = {**y**_1_, …, **y**_*N*_ }from distributions *P* and *Q* respectively, with **x**_*i*_, **y**_*i*_ *∈* ℝ^*d*^, an unbiased MMD estimator’s computational complexity is *𝒪* (*N* ^2^*d*). To mitigate this computational demand, particularly in large single-cell datasets, we propose aligning the transformed sample distributions *μ*_*P*_ (*·*)*∈ ℋ* _*K*_ with the landmark cells *𝒮* from the cell sketching step (see Section 2.1). This gives a sample embedding vector over the *s* landmarks as,

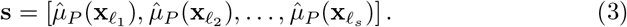

Here, **s** is calculated by evaluating the kernel estimator in (2) with respect to each landmark cell. Moreover, for each of the *m* samples, we can therefore compute its embedding vector. Ultimately, the per-sample embedding vectors are concatenated to form a sample-by-landmark embedding matrix: **S** (see Figure 1**e**), with each row representing a sample embedding vector. Here, 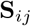 is calculated as the average of the kernel function applied on cells of sample *i* and landmark *𝓁* _*j*_,

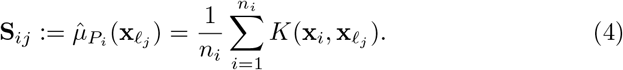

## 3 Experimental Setup

### 3.1 Datasets and pre-processing

We evaluated scLKME’s performance on three single-cell datasets (both CyTOF and single-cell RNA-sequencing (scRNA-seq)) with per-sample outcomes.

#### Multi-species CyTOF data

This CyTOF dataset collects blood samples across five species, including, African Green Monkey (AGM), Cynomolgus Monkey (Cyno), Rhesus Macaque (Rhesus), Human, and Mouse [35]. All samples were profiled with a common antibody panel. We downloaded unstimulated samples (Flow Repository accession id: FR-FCM-Z2Z7), and applied a standard *arcsinh* transformation with a cofactor of 5 to all 24 phenotypic channels of live cells. In our analyses, we focused only on the basal (unstimulated) cells. We sought to generate sample embeddings that captured between-species variation.

#### Preterm CyTOF data

The preterm CyTOF data collected blood samples from 40 donors throughout pregnancy, ranging from 25 weeks to term [36] (data shared by authors.) Each sample was labeled according to its gestational age at birth, including preterm, late preterm, and term. Live cells from unstimulated samples were extracted and each of the 38 channels was transformed using an *arcsinh* transformation with a cofactor of 5. We sought to generate sample embeddings that separated by gestational age.

#### Myocardial infarction scRNA-seq data

The myocardial infarction scRNAseq dataset consists of 115,517 cells from 20 samples, of which 13 were sourced from healthy heart tissues (control) and the remaining 7 were from ischemic zones (IZ) in the heart [20].

### 3.2 Baselines and Evaluation Metrics

scLKME was benchmarked against several standard methods for generating per-sample embeddings, including PhEMD [19], CKME [21], PILOT [20], and Pseudo Bulk [37, 38]. Phenotypic Earth Mover’s Distance (PhEMD) [19] embeds data points from various patient single-cell data by clustering cells into subpopulations and pinpointing cellular variation between subpopulations using earth mover’s distance. Cell Kernel Mean Embedding (CKME) [21] uses randomized Fourier features (RFF) [39] to estimate a predefined kernel between cells and generates sample embeddings using a patient-wise mean pooling on the RFF features. PatIent Level analysis with Optimal Transport (PILOT) [20] detects patient-level distance in single-cell data by estimating sample distribution across predefined cell types using earth mover’s distance. Lastly, we incorporated Pseudo Bulk (PsuBlk) as a baseline, which simply computes an aggregate value for each measured feature across all cells in the sample, using a pooling operation. In our experiments, we used median pooling, where the median of each gene or protein feature was computed across cells, due to its robustness to outliers.

#### Silhouette score

To assess the extent to which the sample embeddings **S** align with their associated labels, we utilized the silhouette score [40], which is defined as:

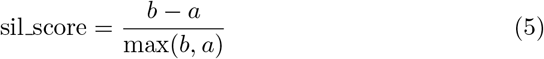

Here, *a* is the mean distance (in the computed embedding space) between a sample and other samples within its class, while *b* denotes the mean distance from the sample to the nearest other samples from a different cluster. We took the average of the silhouette scores across all samples as the metric for evaluating the performance of sample representations. The silhouette score ranges between -1 and 1, with a higher value indicating that the computed sample embeddings are well aligned with sample labels. We employed the `sklearn.metrics.silhouette_score` function in scikit-learn v1.2.2 to compute this score. Notably, as the outputs of PhEMD and PILOT are distance matrices, the inferred distances between samples were used as input for silhouette score computation.

#### Unsupervised clustering

To determine the capability of different methods to retain between-sample structures linked to sample phenotypes and labels, we conducted unsupervised clustering on a *k*-nearest neighbor graph (kNN) graph constructed between samples using the Leiden algorithm [5]. The resolution parameter controlling the ultimate number of clusters under the Leiden algorithm was chosen based on the ground-truth number of sample phenotypes. For PhEMD and PILOT, the output distance matrices were directly employed to construct the between-sample kNN graphs. For other methods, the kNN graph of samples was constructed by computing the cosine similarity between the pairs of samples based on their computed embeddings.

To evaluate the alignment between clusters and sample phenotypes, we used the adjusted rand index (ARI) [41], which is defined as:

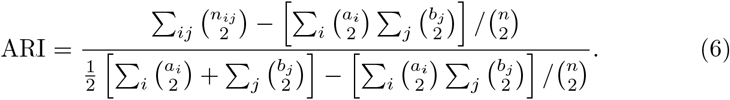

Here, *n*_*ij*_ denotes the count of samples common to both the true cluster (phenotype label) *i* and predicted cluster *j*, while *a*_*i*_ and *b*_*j*_ represent the number of samples in true cluster *i* and predicted cluster *j*, respectively. The ARI ranges from -1 to 1. A score of 0 indicates random labeling, whereas a score of 1 suggests a perfect match. This ARI metric was calculated using the `sklearn.metrics.adjusted_rand_score` function in scikit-learn v1.2.2.

## 4 Experimental Results

### 4.1 Performance for capturing between-sample variation in multi-sample single-cell data

We first evaluated scLKME on the three multi-sample single-cell datasets with known per-sample phenotype labels (section 3.1). This was to verify if scLKME could accurately uncover the true patterns of inter-sample variation based on the computed embeddings. As shown in Table 1, our scLKME method achieved the best alignment between computed embeddings and per-sample phenotypes on both the Preterm CyTOF dataset and the Myocardial infarction scRNA-seq dataset, with respective ARI scores of 0.227 and 0.198, and silhouette scores of 1.000 and 0.720. On the Multi-species dataset, scLKME presented a comparable performance with CKME, the leading method in terms of ARI score. Notably, CKME showed significantly poor results on both the Preterm and the Myocardial infarction datasets. Given that CKME relied on a mean pooling operation to encode samples, we suspect this is due to the information loss in the pooling step, especially in the presence of cell heterogeneities. Pseudo Bulk, another pooling-based approach, consistently gave fairly good results by employing median estimations on cell embeddings per sample. Although this pooling strategy worked well when cell embeddings exhibited unimodal distribution, these sample embeddings would be skewed in cases of multi-modality. PILOT’s performance was marked as `NA` in the two CyTOF datasets because PILOT requires using ground-truth cell-type labels, which were not available and shows its limitation as an automated tool. Second to scLKME, PhEMD achieved strong performance based on silhouette and ARI scores on the Multi-species and the Myocardial infarction datasets respectively.

**Table 1.** Performance of the alignment between computed sample embeddings and sample phenotype labels.

| Datasets | Metric | Method |  |  |  |  |
| --- | --- | --- | --- | --- | --- | --- |
|  |  | PhEMD[19] | PILOT[20] | Pseudo Bulk | CKME[21] | scLKME(Ours) |
| Multi-species | ARI | 0.621 | NA | 0.608 | **0.737** | 0.730 |
|  | Sil_score | **0.347** | NA | 0.271 | 0.136 | 0.260 |
| Preterm | ARI | 0.223 | NA | **0.227** | 0.037 | **0.227** |
|  | Sil_score | 0.106 | NA | 0.171 | 0.047 | **0.198** |
| Myoc. Infarc. | ARI | **1.000** | 0.799 | **1.000** | -0.031 | **1.000** |
|  | Sil_score | 0.706 | 0.492 | 0.497 | 0.007 | **0.720** |

#### Robustness and runtime efficiency

We next assessed various methods based on hyperparameter robustness and computational efficiency. In terms of robustness, Pseudo Bulk and PILOT emerged superior since their outputs are deterministic. When it comes to runtime efficiency, Pseudo Bulk, PILOT and CKME are the most efficient methods because they only use linear- or pooling-based calculations. For PhEMD, its efficiency is limited by the algorithm used in the clustering step.

For robustness and efficiency, we further compared our method with PhEMD. We evaluated how the number of landmarks affected performance, varying this parameter as *s* = 64, 128, 256, 512. Similarly, PhEMD was evaluated by using varying number of clusters as `n_phate_cluster` = 8, 16, 24, 32. The results are shown in Figure 2a. Compared with PhEMD, our method demonstrated significantly reduced variance in terms of both ARI and silhouette scores. Figure 2b represents the runtime comparison of PhEMD and scLKME under the varying hyperparameters above. Due to PhEMD’s limited scalability, its runtime was measured on downsampled versions of the Multi-species and Preterm CyTOF datasets, each containing 100,000 total cells. In contrast, scLKME was tested on the entire dataset. Despite this, our method still surpassed PhEMD in speed significantly, suggesting that scLKME is more applicable in large singlecell datasets.

**Fig. 2.**
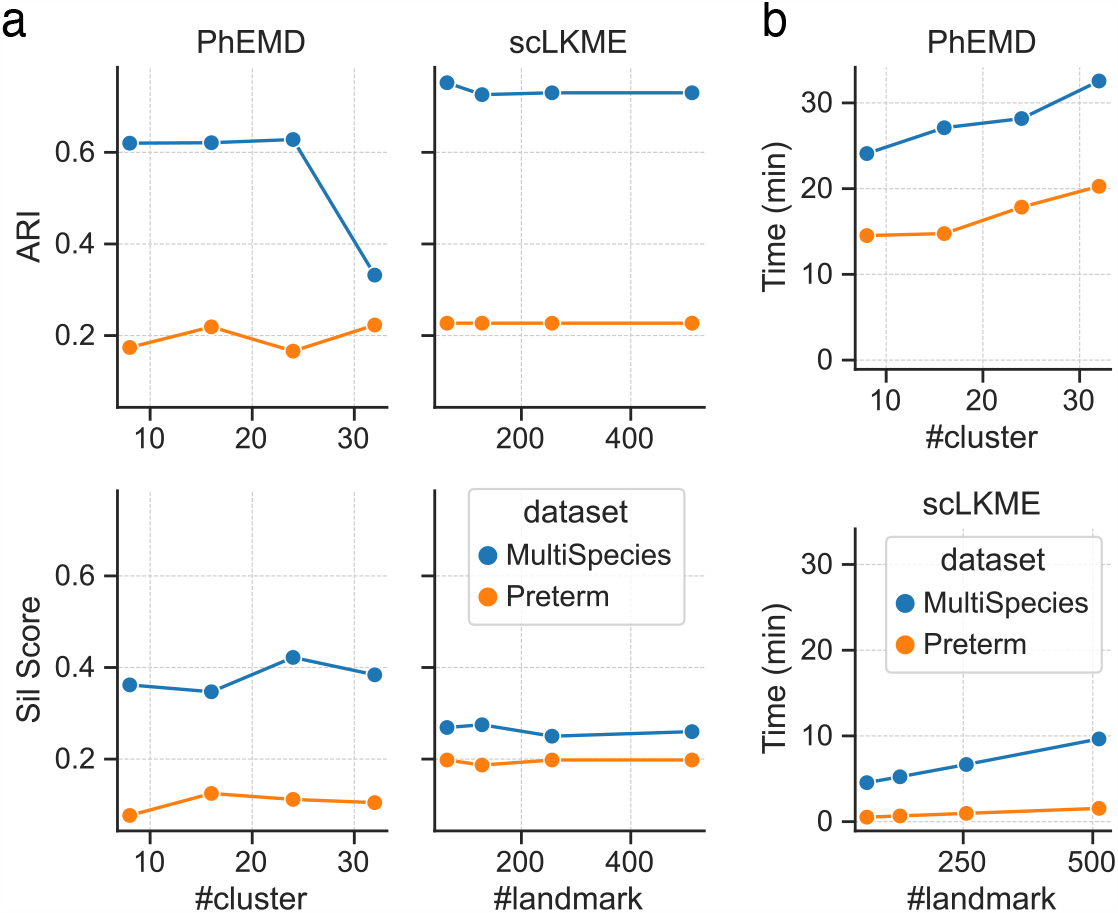
**a**. Performance comparison between PhEMD and scLKME on the Multi-species and Preterm CyTOF datasets across various hyperparameters: PhEMD varied the number of clusters 8, 16, 24, 32, and scLKME varied the number of landmark cells: 64, 128, 256, 512. **b**. Corresponding runtime analysis for PhEMD and scLKME using the the hyperparameters in **a**.

### 4.2 Sample-level unsupervised analysis using scLKME in single-cell cohorts

To illustrate scLKME’s capacity for unsupervised sample-level analysis, we applied it to the Preterm CyTOF dataset introduced in Ref. [36]. Cells across all 40 patients were visualized with a two-dimensional UMAP embedding (Figure 3a) and colored by the sample-level labels of either *Preterm, Late Preterm*, or *Term* birth. In Figure 3b, scLKME identified a total of 512 landmark cells, effectively capturing the overall cellular landscape visualized in Figure 3a. All profiled samples were embedded in two-dimensions with PHATE [9] in Figure 3c and showed strong organization by gestational age along the curve, with samples transitioning from Term to Late Preterm and Preterm with reducing gestational age. This suggests that scLKME can effectively capture the key phenotypic variations across samples. Applying dimension reduction on the sample embeddings via diffusion maps [6] further validated our results. The first non-trivial diffusion map component (DC1) distinctively separated sample categories (p-value *<* 0.01, Figure 3d) and correlated strongly with gestational age (p-value = 1e-9, Appendix Figure 4).

**Fig. 3.**
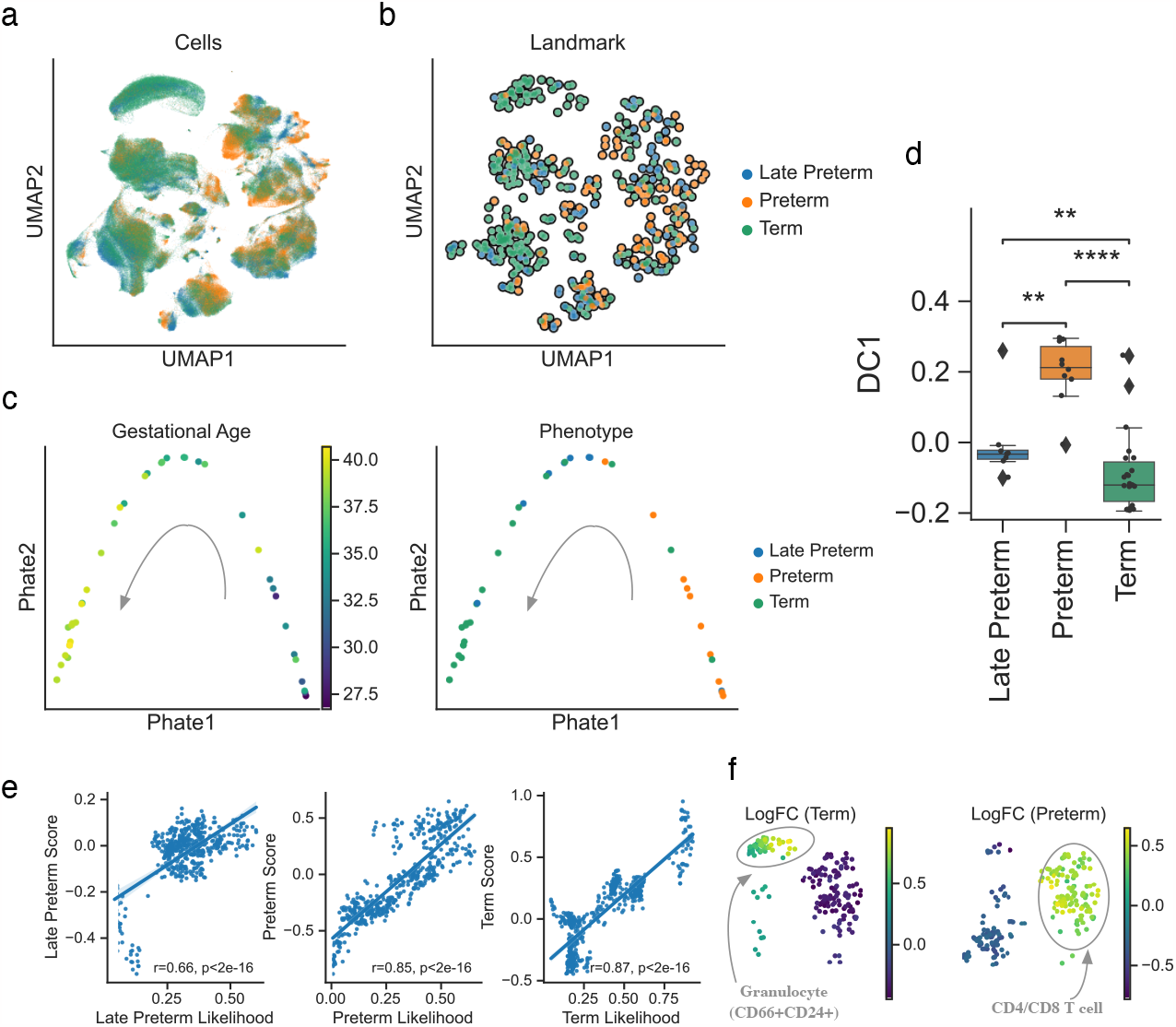
Unsupervised analysis of the Preterm CyTOF dataset. **a**. UMAP visualizations of cells across samples colored by phenotype. **b**. Selected landmark cells in the sketching step. **c**. PHATE visualization of scLKME sample embeddings, colored by gestational age (left) and phenotype (right). **d**. Distribution of the first non-trivial component of diffusion map for sample embeddings. Scatter plot of the second diffusion component vs. gestational age. **e**. Scatter plot of MELD likelihood vs. phenotype importance score from the Wilcoxon rank sum test on the landmarks. **f**. Statistically significant (FDR *<* 1e-3) landmark cells with for Term and Preterm phenotypes, colored by log fold change (LogFC) values obtained by one vs. rest testing of term (left) and preterm (right) landmarks respectively against all other landmarks. The analysis highlights granulocytes as being characteristic of term birth and both CD4 and CD8 T-cells being prototypical of preterm birth.

**Fig. 4.**
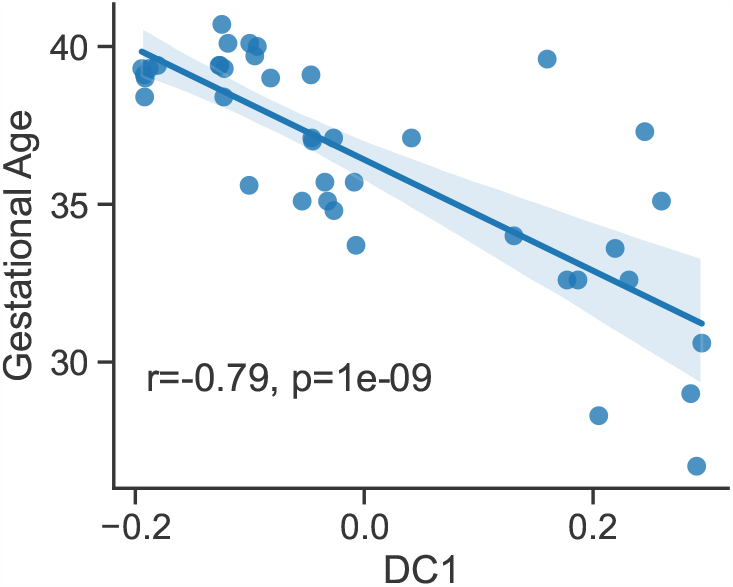
Scatter plot of the first non-trivial component of diffusion map (DC1) vs. gestational age. The correlation between DC1 and gestational age is 0.79 with p-value = 1e-9.

Recall that scLKME computes sample embeddings by averaging kernel affinities across cells for each landmark (Eq. 4). Thus, scLKME’s sample embeddings can be viewed as “soft” abundance scores, which is similar to the idea behind PILOT and PhEMD but excluding the manual or automatic cell clustering step. To verify this, we applied one-vs-rest Wilcoxon rank-sum testing to the sample embeddings, with the log fold change (LogFC) denoting the enrichment score for each phenotype label. As a reference of comparison, we used MELD [42] to compute landmark likelihood scores for each phenotype label. Their strong correlation (p-value *<* 2e-16, Figure 3e) demonstrated the potential of our approach to summarize sample variation based on abundances. This is an extension to PILOT and PhEMD since scLKME doesn’t need clustering or manual annotation to identify cell types. Significant landmarks were visualized in Figure 3f (FDR *<* 1e-3), revealing that patients with Term and Preterm labels were differentially enriched in distinct cell populations. The Preterm patients had higher proportions of T cells overall (both CD8^+^ and CD4^+^), whereas Term patients exhibited increased antigen presentation cells (granulocytes (CD66^+^CD24^+^) (Figure 3f), suggesting the presence of antigen- and cytokine-specific immune cell signaling. These results aligned with the finds identified through manual gating in Ref. [36].

## 5 Conclusion

In this work, we proposed scLKME, an unsupervised landmark-based multicellular sample embedding method for adequately encoding phenotype-associated patterns of cellular variation in multi-sample single-cell cohorts. We validated our approach by applying it to three single-cell datasets, including both scRNAseq and CyTOF datasets, and compared it with other recently-developed sample embedding methods. Our results demonstrated scLKME can better capture variation in cellular heterogeneity across samples by mapping their distributions into RKHS through kernel mean embeddings. In addition, we further showed that scLKME was robust to the number of landmark cells and efficient enough for large single-cell datasets with millions of cells. In conclusion, our results has demonstrated scLKME can be effectively applied for generating sample embeddings and visualizing them in multi-sample single-cell data. In future work, we plan to extend scLKME to multi-modal single data, where the objective is to generate sample embeddings across modalities.

## Acknowledgements

This work is supported by The National Institutes of Allergy and Infectious Diseases of the National Institutes of Health under the award number 1R21AI171745-01A1 (NS). We would like to thank Dr. Jolene Ranek for her insightful discussions on this work.

## Notes

### Competing Interest Statement

The authors have declared no competing interest.

## Reference

[1] Vincent D Blondel et al. “Fast unfolding of communities in large networks”. In: Journal of statistical mechanics: theory and experiment 2008.10 (2008), P10008.

[2] Jacob H Levine et al. “Data-driven phenotypic dissection of AML reveals progenitor-like cells that correlate with prognosis”. In: Cell 162.1 (2015), pp. 184–197.

[3] Vladimir Yu Kiselev et al. “SC3: consensus clustering of single-cell RNA-seq data”. In: Nature methods 14.5 (2017), pp. 483–486.

[4] Peijie Lin, Michael Troup, and Joshua WK Ho. “CIDR: Ultrafast and accurate clustering through imputation for single-cell RNA-seq data”. In: Genome biology 18.1 (2017), pp. 1–11.

[5] Vincent A Traag, Ludo Waltman, and Nees Jan Van Eck. “From Louvain to Leiden: guaranteeing well-connected communities”. In: Scientific reports 9.1 (2019), p. 5233.

[6] Ronald R Coifman and Stéphane Lafon. “Diffusion maps”. In: Applied and computational harmonic analysis 21.1 (2006), pp. 5–30.

[7] Laurens Van der Maaten and Geoffrey Hinton. “Visualizing data using t-SNE.” In: Journal of machine learning research 9.11 (2008).

[8] Bo Wang et al. “Visualization and analysis of single-cell RNA-seq data by kernel-based similarity learning”. In: Nature methods 14.4 (2017), pp. 414–416.

[9] Kevin R Moon et al. “Visualizing structure and transitions in high-dimensional biological data”. In: Nature biotechnology 37.12 (2019), pp. 1482–1492.

[10] Etienne Becht et al. “Dimensionality reduction for visualizing single-cell data using UMAP”. In: Nature biotechnology 37.1 (2019), pp. 38–44.

[11] Zhicheng Ji and Hongkai Ji. “TSCAN: Pseudo-time reconstruction and evaluation in single-cell RNA-seq analysis”. In: Nucleic acids research 44.13 (2016), e117–e117.

[12] Laleh Haghverdi et al. “Diffusion pseudotime robustly reconstructs lineage branching”. In: Nature methods 13.10 (2016), pp. 845–848.

[13] Xiaojie Qiu et al. “Reversed graph embedding resolves complex single-cell trajectories”. In: Nature methods 14.10 (2017), pp. 979–982.

[14] Kelly Street et al. “Slingshot: cell lineage and pseudotime inference for single-cell transcriptomics”. In: BMC genomics 19 (2018), pp. 1–16.

[15] FAlexander Wolf et al. “PAGA: graph abstraction reconciles clustering with trajectory inference through a topology preserving map of single cells”. In: Genome biology 20 (2019), pp. 1–9.

[16] Eirini Arvaniti and Manfred Claassen. “Sensitive detection of rare disease-associated cell subsets via representation learning”. In: Nature communications 8.1 (2017), pp. 1–10.

[17] Lukas M Weber et al. “diffcyt: Differential discovery in high-dimensional cytometry via high-resolution clustering”. In: Communications biology 2.1 (2019), p. 183.

[18] Haidong Yi and Natalie Stanley. “CytoSet: Predicting clinical outcomes via set-modeling of cytometry data”. In: Proceedings of the 12th ACM Conference on Bioinformatics, Computational Biology, and Health Informatics. 2021, pp. 1–8.

[19] William S Chen et al. “Uncovering axes of variation among single-cell cancer specimens”. In: Nature methods 17.3 (2020), pp. 302–310.

[20] Mehdi Joodaki et al. “Detection of PatIent-Level distances from single cell genomics and pathomics data with Optimal Transport (PILOT)”. In: bioRxiv (2022). doi: 10.1101/2022.12.16.520739. eprint: https://www.biorxiv.org/content/early/2022/12/19/2022.12.16.520739.full.pdf. url: https://www.biorxiv.org/content/early/2022/12/19/2022.12.16.520739.

[21] Siyuan Shan et al. “Transparent single-cell set classification with kernel mean embeddings”. In: Proceedings of the 13th ACM International Conference on Bioinformatics, Computational Biology and Health Informatics. 2022, pp. 1–10.

[22] Brian Hie et al. “Geometric sketching compactly summarizes the single-cell transcriptomic landscape”. In: Cell systems 8.6 (2019), pp. 483–493.

[23] Yutian Chen, Max Welling, and Alex Smola. “Super-Samples from Kernel Herding”. In: UAI’10. Arlington, Virginia, USA: AUAI Press, 2010, p. 109–116. isbn: 9780974903965.

[24] Vishal Athreya Baskaran et al. “Distribution-based sketching of singlecell samples”. In: Proceedings of the 13th ACM International Conference on Bioinformatics, Computational Biology and Health Informatics. 2022, p. 1–10.

[25] Nachman Aronszajn. “Theory of reproducing kernels”. In: Transactions of the American mathematical society 68.3 (1950), pp. 337–404.

[26] Kenji Fukumizu, Francis R Bach, and Michael I Jordan. “Dimensionality reduction for supervised learning with reproducing kernel Hilbert spaces”. In: Journal of Machine Learning Research 5.Jan (2004), pp. 73–99.

[27] Bharath K Sriperumbudur, Kenji Fukumizu, and Gert RG Lanckriet. “Universality, Characteristic Kernels and RKHS Embedding of Measures.” In: Journal of Machine Learning Research 12.7 (2011).

[28] Alain Berlinet and Christine Thomas-Agnan. Reproducing kernel Hilbert spaces in probability and statistics. Springer Science & Business Media, 2011.

[29] Alex Smola et al. “A Hilbert space embedding for distributions”. In: Algorithmic Learning Theory: 18th International Conference, ALT 2007, Sendai, Japan, October 1–4, 2007. Proceedings 18. Springer. 2007, p. 13–31.

[30] Alfred Müller. “Integral probability metrics and their generating classes of functions”. In: Advances in applied probability 29.2 (1997), pp. 429–443.

[31] Karsten M Borgwardt et al. “Integrating structured biological data by kernel maximum mean discrepancy”. In: Bioinformatics 22.14 (2006), e49–e57.

[32] Arthur Gretton et al. “A kernel two-sample test”. In: The Journal of Machine Learning Research 13.1 (2012), pp. 723–773.

[33] Yujia Li, Kevin Swersky, and Rich Zemel. “Generative moment matching networks”. In: International conference on machine learning. PMLR. 2015, p. 1718–1727.

[34] Gintare Karolina Dziugaite, Daniel M. Roy, and Zoubin Ghahramani. “Training generative neural networks via Maximum Mean Discrepancy optimization”. In: Proceedings of the Thirty-First Conference on Uncertainty in Artificial Intelligence, UAI 2015, July 12-16, 2015, Amsterdam, The Netherlands. Ed. by Marina Meila and Tom Heskes. AUAI Press, 2015, p. 258–267. url: http://auai.org/uai2015/proceedings/papers/230.pdf.

[35] Zachary B Bjornson-Hooper et al. “A comprehensive atlas of immunological differences between humans, mice, and non-human primates”. In: Frontiers in immunology 13 (2022).

[36] Laura S. Peterson et al. “Single-Cell Analysis of the Neonatal Immune System Across the Gestational Age Continuum”. In: Frontiers in Immunology 12 (Aug. 2021), p. 714090. doi: 10.3389/fimmu.2021.714090.

[37] Mark D Robinson, Davis J McCarthy, and Gordon K Smyth. “edgeR: a Bioconductor package for differential expression analysis of digital gene expression data”. In: bioinformatics 26.1 (2010), pp. 139–140.

[38] Michael I Love, Wolfgang Huber, and Simon Anders. “Moderated estimation of fold change and dispersion for RNA-seq data with DESeq2”. In: Genome biology 15.12 (2014), pp. 1–21.

[39] Ali Rahimi and Benjamin Recht. “Random features for large-scale kernel machines”. In: Advances in neural information processing systems 20 (2007).

[40] Peter J Rousseeuw. “Silhouettes: a graphical aid to the interpretation and validation of cluster analysis”. In: Journal of computational and applied mathematics 20 (1987), pp. 53–65.

[41] Lawrence Hubert and Phipps Arabie. “Comparing partitions”. In: Journal of classification 2 (1985), pp. 193–218.

[42] Daniel B Burkhardt et al. “Quantifying the effect of experimental perturbations at single-cell resolution”. In: Nature biotechnology 39.5 (2021), pp. 619–629.

